# Recovery of protein synthesis to measure transcription-coupled DNA repair in living cells and tissues

**DOI:** 10.1101/2023.01.24.525327

**Authors:** Melanie van der Woude, Carlota Davó-Martínez, Karen L. Thijssen, Wim Vermeulen, Hannes Lans

## Abstract

Transcription-coupled nucleotide excision repair (TC-NER) is an important DNA repair mechanism that protects against the negative effects of transcription-blocking DNA lesions. Hereditary TC-NER deficiency causes pleiotropic and often severe neurodegenerative and progeroid symptoms. Multiple assays have been developed for the clinic and for research to measure TC-NER activity, which is hampered by the relatively low abundance of repair events taking place in transcribed DNA. ‘Recovery of RNA Synthesis’ is widely used as indirect TC-NER assay based on the notion that lesion-blocked transcription only resumes after successful TC-NER. Here, we show that measuring novel synthesis of a protein that has been degraded prior to DNA damage induction is an equally effective but more versatile manner to indirectly monitor TC-NER. This ‘Recovery of Protein Synthesis’ (RPS) assay is readily adaptable for use with different degradable proteins and readouts, including fluorescence imaging and immunoblot. Moreover, with the RPS assay TC-NER activity can be measured in real-time, in various living cells types and even in differentiated tissues of living organisms. As example, we show that TC-NER capacity declines in aging muscle tissue of *C. elegans*. Therefore, the RPS assay constitutes an important novel clinical and research tool to investigate transcription-coupled DNA repair.

## INTRODUCTION

DNA damage induced by various environmental and metabolism-derived agents continuously threatens the integrity and functionality of DNA. DNA damage interferes with essential DNA-transacting processes like transcription and replication and contributes to aging and causes mutagenesis, leading to cancer (1). Various dedicated DNA repair pathways maintain genomic integrity by removing DNA damage depending on the type of lesion, its genomic location and the cell cycle stage. Nucleotide excision repair (NER) is an important DNA repair mechanism that protects organisms against cancer and aging by removing different DNA helix-distorting lesions, such as those induced by UV light, cancer therapeutics like cisplatin, metabolism-derived aldehydes and various genotoxic environmental chemicals including polycyclic aromatic hydrocarbon and aromatic amines found in smoke and cooked food (2, 3).

NER is initiated by lesion detection via global genome NER (GG-NER) anywhere in the entire genome, or via transcription-coupled NER (TC-NER) exclusively in the transcribed strands of active genes. Lesion detection via GG-NER is mediated by the XPC protein, which continuously probes DNA and recognizes lesions by interacting with the DNA strand opposite of the lesion and inserting a β-hairpin domain into the distorted DNA duplex (4–6). XPC exists in complex with CETN2 and RAD23B and is aided by the CRL4^DDB2^ ubiquitin ligase complex when lesions are difficult to detect and/or when chromatin needs to be re-organized (7, 8). Lesion detection via TC-NER occurs when lesions block forward progression of elongating RNA polymerase II (Pol II), which triggers the stable association of the CSB protein with Pol II (9). CSB then recruits the CRL4^CSA^ ubiquitin ligase complex and additional TC-NER factors (10). Lesion detection via either XPC or CSB leads to sequential recruitment of core NER factors, including TFIIH and XPA, which unwind DNA and check for the presence of damage, and the endonucleases ERCC1-XPF and XPG that excise a 22-30 bp DNA stretch containing the lesion (11, 12). The resulting gap is then filled in by DNA synthesis mediated by replication factors, DNA polymerases and ligases.

The biological significance of NER is illustrated by the severe cancer-prone, developmental and progeroid symptoms associated with hereditary mutations in NER genes (13). GG-NER deficiency causes xeroderma pigmentosum (XP), which manifests as photosensitive skin and strong cancer predisposition (14). Mutations in TC-NER factors cause either the mild UV-sensitive syndrome (UV^S^S) or the much more severe Cockayne syndrome (CS) (15, 16). Mutations in genes involved in both GG-NER and TC-NER can cause severe XP, often also combined with CS features, or a photosensitive form of trichothiodystrophy (TTD) (17, 18). UV^S^S is mainly characterized by telangiectasia and sun sensitivity of the skin, whereas CS is characterized by a pleiotropic range of severe symptoms including growth failure, progressive organ decline and neurodegeneration and segmental progeria. It is currently not understood why different mutations in TC-NER factors lead to this wide array of symptoms, but it is hypothesized that this may be related to differences in clearance of lesion-stalled Pol II (19). TTD is characterized by brittle hair and nails, ichthyosis and progressive mental and physical retardation and is thought to be mainly caused by problems with gene expression (20). Interestingly, research on progeroid features of TC-NER deficiency disorders, in humans and mouse models, has revealed that accumulating DNA damage interfering with transcription is one of the major underlying causes of aging (21, 22). To study the etiology of aging, and how this affects organs differently, it is therefore important to be able to investigate TC-NER activity in different types of cells *in vivo*.

Both the clinical diagnosis of NER disorders as well as the discovery of novel genes involved in NER depends heavily on the availability of reliable and straightforward assays that can discriminate GG-NER and TC-NER deficiency. GG-NER activity can be determined by measuring unscheduled DNA synthesis (UDS) that restores the single-stranded DNA gap resulting from DNA damage excision. Originally, this assay was used to link DNA repair defects to XP and was based on GG-NER-dependent incorporation of radioactively labeled thymine analogs (23). Nowadays, these have been replaced by 5-ethynyl-2’-deoxyuridine, which can be visualized by fluorescent labeling (24, 25). TC-NER activity is more difficult to detect largely due to the fact that this repair pathway only removes a minority of lesions, i.e., only those that block transcription. Originally, TC-NER was demonstrated using a Southern blot-based assay that showed preferential repair in the transcribed strands of active genes (26), which was subsequently used to show TC-NER deficiency in cells from CS patients (27). Since then, multiple assays have been developed to monitor TC-NER activity either directly, such as the gene-specific qPCR (28), comet-FISH (29), the amplified UDS (30) and strand-specific ChIP-Seq (31) assays, or indirectly, such as the host cell reactivation (32) and the Recovery of RNA synthesis (RRS) assays. Because of its ease of use, the RRS assay is commonly used in NER research and in the clinic. It is based on the notion that DNA damage inhibits transcription and that this can only resume if TC-NER is active and removes the damage (33). Transcription recovery can be measured by labeling RNA with radioactive or bromo-uridine or, as is nowadays more common, 5-ethynyluridine that can be fluorescently labeled (24, 25). A drawback of RRS and most other techniques is that these assays only work in cells in culture and cannot easily be used to measure repair in real-time or *in vivo*.

Here, we show the utility of a novel versatile assay to indirectly monitor TC-NER based on the idea that not only transcription but also translation will only resume after DNA damage induction if TC-NER is proficient. We show that this ‘recovery of protein synthesis’ (RPS) assay is robust and reliable and can be performed by monitoring the novel synthesis of different proteins, using both fluorescence imaging as well as immunoblot as readout. Moreover, the RPS assay can be used to measure TC-NER activity in real-time both in living cells in culture as well as *in vivo* in differentiated cell types of a living organism.

## MATERIAL AND METHODS

### Cell culture and treatment conditions

U2OS (ATCC) and Hep3B cells stably expressing EGFP-AR (34, 35) were cultured in DMEM (Invitrogen), supplemented with 10% fetal bovine serum (FBS; Bodinco BV), and 1% penicillin/streptomycin (Sigma). C5RO-SV40 (36), CSBE-SV40 (GM01856; Coriell Institute) and CS1AN-SV40 (37) fibroblasts were cultured in HAM’s F10 (Invitrogen) supplemented with 15% FBS and 1% penicillin/streptomycin. Transient siRNA-mediated knockdown was achieved using transfection with Lipofectamine RNAiMAX (Invitrogen) according to the manufacturer’s instruction. The siRNA oligonucleotides (Horizon Discovery) used were: CTRL 5’-UGGUUUACAUGUUGUGUGA-3’, CSB 5’-GCAUGUGUCUUACGAGAUA-3’, XPF 5’-AAGACGAGCUCACGAGUAU-3’, and XPC 5’-GCAAAUGGCUUCUAUCGAA-3’. Knockdown was confirmed by immunoblot (Supplementary Figure 1A). Cells were incubated with 50 nM dTAG-13 (Tocris) to degrade EGFP-FKBP^F36V^, 100 nM ARV-110 (Selleckchem) to degrade EGFP-AR and 250 nM PROTAC FAK degrader 1 (MedChemExpress) to degrade PTK2/FAK, as indicated. XPB depletion was achieved by 4 h spironolactone (Sigma) incubation with the indicated doses. For induction of DNA damage by UV, cells were exposed to 6 J/m^2^ UV-C light (254 nm; TUV lamp, Phillips). For cisplatin induction of DNA damage, cells were exposed for 2 h to 100 µM cisplatin (Sigma).

### Plasmid and cell line generation

To generate U2OS cells stably expressing EGFP-FKBP^F36V^, a gene fragment containing *EGFP*-*FKBP*^*F36V*^ with nuclear localization signal behind a PGK promoter and *BLAST* behind an EF1a promoter, flanked by *AAVS1* homology sequences for homology directed repair, was generated by Integrated DNA technologies. The gene fragment was cloned into pUC57 and transfected into U2OS cells. After selection with blasticidin, a clonal cell line stably expressing EGFP-FKBP^F36V^ was isolated.

### Live-cell confocal laser-scanning microscopy

For live cell imaging, cells were grown on 24-mm coverslips and imaged using a Leica SP5 confocal microscope equipped with an environmental chamber set to 37°C and 5% CO2. Confocal images were recorded every hour after UV irradiation, as indicated. Data collection and analysis was performed using LAS X software (Leica).

### Fluorescence imaging

For imaging of fluorescence levels, cells were grown on 24-mm coverslips and fixed using 3.6% formaldehyde (Sigma) diluted in PBS. 4′,6-diamidino-2-phenylindole (DAPI; Sigma) staining was performed in PBS for 15 min at room temperature. Subsequently, coverslips were washed twice with PBS supplemented with 0.1 % Triton-X-100 and once with PBS and mounted with aqua-poly/mount (Polysciences Inc). Cells were imaged using a Zeiss LSM700 microscope equipped with a Plan-Apochromat 40x/1.3 Oli DIC M27 immersion lens (Carl Zeiss Microimaging Inc.). EGFP-FKBP^F36V^ or EGFP-Androgen receptor expression in the nucleus was quantified in Fiji ImageJ.

### Immunoblot and antibodies

For immunoblotting, cells were lysed in sample buffer (0.125 M Tris-HCl pH 6.8, 2% SDS, 0.005% bromophenol blue, 21% glycerol, 4% β-mercaptoethanol) and boiled for 5 min at 98°C. Lysates were separated by sodium dodecyl sulphate-polyacrylamide gel electrophoresis (SDS–PAGE) using NuPAGE™ 4-12% Bis-Tris gels (Invitrogen) and transferred to Polyvinylidene difluoride (PVDF) membrane (0.45 µm) (Millipore). Membranes were blocked in 5% BSA (Sigma) for 30 min at room temperature and incubated for 2 h at room temperature or overnight at 4°C with primary antibodies against GFP (Roche), androgen receptor (Invitrogen), PTK2 (Invitrogen), XPB (Abcam), CSB (Bethyl), XPF (Santa Cruz), XPC (Bethyl), Tubulin (Sigma Aldrich) or Ku70 (Santa Cruz). Membranes were incubated with secondary antibodies coupled to IRDyes (Sigma) and scanned using an Odyssey CLx infrared scanner (LiCor).

### *C. elegans* strains and experimental handling

*C. elegans* were cultured according to standard methods (38). Strains and alleles used were CA1202 (*eSi57[P(eft-3)::TIR1::mRuby] II; ieSi58 [P(eft-3)::AID::GFP] IV*) (39); HAL526 (*ieSi57 [P(eft-3)::TIR1::mRuby] II; ieSi58 [P(eft-3)::AID::GFP] IV; csb-1(ok2335)*); HAL534 (*ieSi57 [P(eft-3)::TIR1::mRuby] II; ieSi58 [P(eft-3)::AID::GFP] IV; xpc-1(tm3886)*); HAL535 (*ieSi57 [P(eft-3)::TIR1::mRuby] II; ieSi58 [P(eft-3)::AID::GFP] IV; csb-1(ok2335); xpc-1(tm3886)*). HAL526, HAL534 and HAL535 strains were generated by crossing CA1202 with *xpc-1(tm3886)* and/or *csb-1(ok2335)* mutants (40) and genotyped by PCR and sequencing. For the recovery of protein synthesis assay, animals of similar age were depleted of AID::GFP expression by culturing for 2 h on NGM plates containing 100 µM auxin (3-indoleacetic acid; Sigma). Directly after depletion, animals were mock-treated or irradiated with 120 J/m^2^ UV-B (Philips TL-12 tubes, 40W) and allowed to recover for 48 h on NMG plates containing OP50 *E. coli* food. AID::GFP levels were measured by imaging fluorescence in body wall muscle cells in the head of living animals on a Leica TCS SP8 microscope (LAS AF software, Leica). AID::GFP expression levels were quantified in Fiji ImageJ.

## RESULTS AND DISCUSSION

### Recovery of protein synthesis after DNA damage depends on transcription-coupled NER

As the RRS assay is based on measuring the ability of cells to recover transcription after DNA damage induction, we reasoned that the ability of cells to re-synthesize a degraded protein could be used as indirect TC-NER readout as well (Figure 1A). After protein degradation, ongoing transcription will produce new proteins, which will still happen if cells incur transcription-blocking DNA damage that is repaired. However, in the absence of TC-NER, transcription blockage will persist and less or no new protein will be produced. We first tested this idea using GFP, as imaging the re-synthesis of a fluorescent reporter will allow the monitoring of TC-NER activity in real-time in living cells. We therefore generated a DNA construct expressing EGFP fused to the FKBP^F36V^ degradation tag (dTAG) (41) and to a nuclear localization signal, which can be efficiently knocked-in at the AAVS1 locus using CRISPR-Cas9 in any cell type (42) and selected for by blasticidin (Figure 1B). Proteins fused to the mutant FKBP^F36V^ dTAG are rapidly degraded by the proteasome via polyubiquitination by the CRL4^Cereblon^ E3 ubiquitin ligase complex, after addition of a heterobifunctional dTAG ligand such as dTAG-13 to cells (41). Rapid protein depletion using this system before DNA damage induction will therefore enable us to determine if, after DNA damage induction, novel protein synthesis and therefore DNA repair takes place. After establishing human osteosarcoma U2OS cells stably expressing EGFP-FKBP^F36V^ in the nucleus, we found by fluorescence imaging of fixed cells that addition of dTAG-13 led to rapid loss of the nuclear fluorescent signal, which re-appeared upon washing away dTAG-13 (Figure 1C).

**Figure 1.**
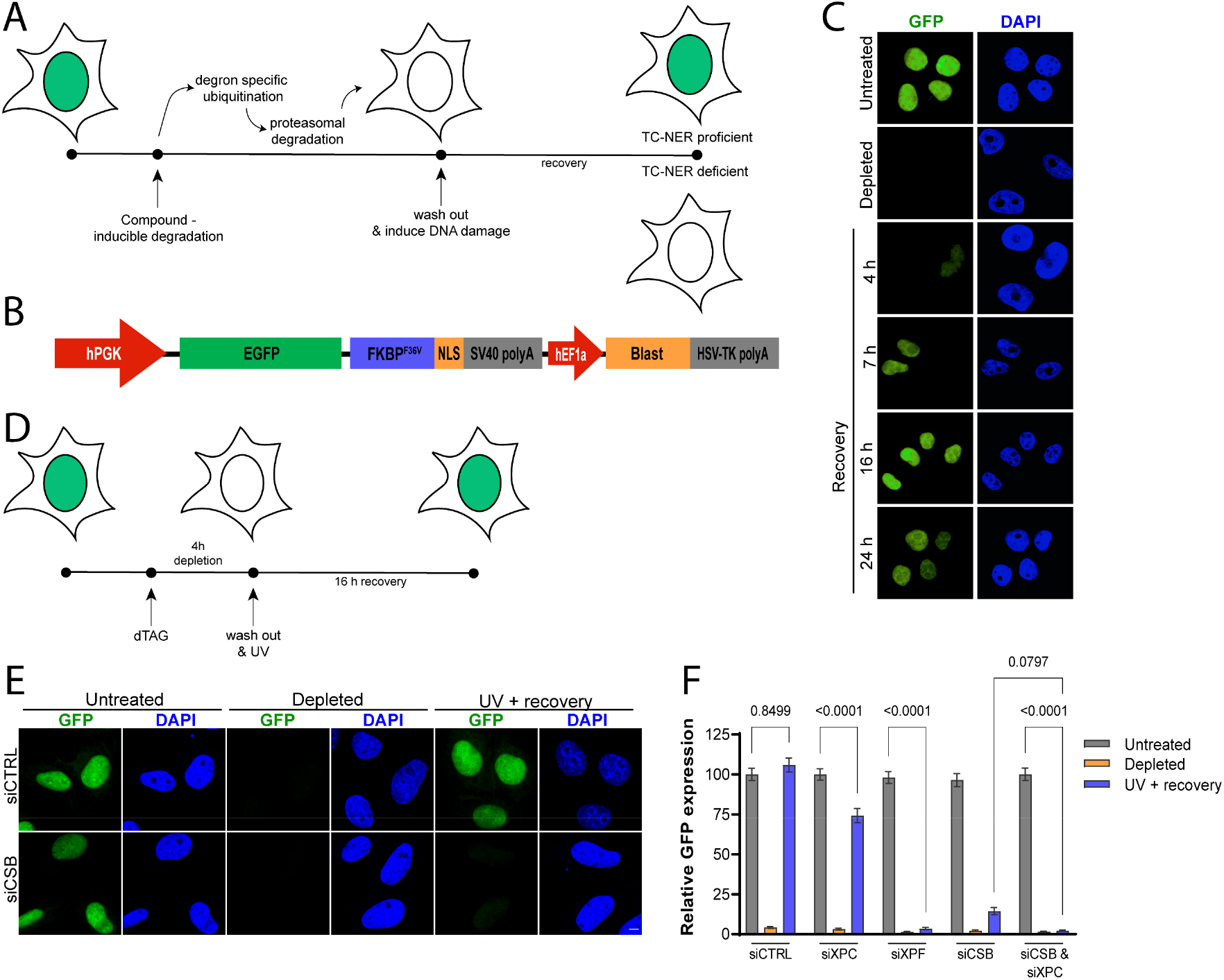
Recovery of protein synthesis after DNA damage depends on transcription-coupled NER. (**A**) Schematic depiction of the rationale of the Recovery of Protein Synthesis (RPS) assay. First a protein (indicated by green shading in the nucleus) is degraded in cells, after which cells are exposed to a DNA damaging agent and the novel synthesis of the protein is monitored in time. The protein will only be produced in TC-NER proficient cells after DNA damage induction. (**B**) Scheme of the transgene knocked-in at the AAVS1 locus to drive EGFP-FKBP^F36V^ from the PGK promoter and blasticidin (for selection) from the hEF1a promoter. (**C**) Representative images of fixed U2OS cells stably expressing EGFP-FKBP^F36V^, before treatment (untreated) and directly (depleted) or several hours (as indicated) after treatment with dTAG13. (**D**) Timing scheme of the RPS assay using UV-C irradiation. (**E**) Representative images of fixed EGFP-FKBP^F36V^-expressing U2OS cells transfected with control (siCtrl) or CSB (siCSB) siRNA that were either untreated, incubated with dTAG13 for 4 h (‘depleted’) or incubated with dTAG13 and irradiated with 6 J/m^2^ UV-C and left to recover for 16 h (‘UV + recovery’). Scale bars 5 µm (**F**) Quantification of GFP expression levels from siRNA-treated cells imaged and treated as explained in (**E**). Mean and S.E.M. of three independent experiments. Statistical difference determined with One way ANOVA is indicated.

Subsequently, we tested if EGFP protein synthesis is dependent on TC-NER after DNA damage induction. We treated the cells with control siRNA or siRNA against the GG- and TC-NER factor XPF, the GG-NER factor XPC and/or the TC-NER factor CSB, after which we depleted EGFP and UV irradiated the cells (Figure 1D). Untreated cells, which were not exposed to dTAG-13 and UV-irradiation, were used for comparison. In control siRNA-treated cells, the EGFP fluorescence levels returned to similar levels as in untreated cells within 16 h (Figure 1E and F). In sharp contrast, EGFP protein synthesis was completely abolished in XPF-depleted cells and almost completely abolished in CSB-depleted cells. Depletion of XPC led to a minor reduction in EGFP levels and further reduced EGFP levels in CSB-depleted cells to the same level as in XPF-depleted cells. Importantly, if cells were not UV irradiated, EGFP fluorescence recovered to similar levels in all siRNA-treated cells after EGFP depletion (Supplementary Figure 1B). These results demonstrate that protein resynthesis after its depletion can be used as readout of TC-NER capacity after DNA damage induction.

Our results suggest that in UV-irradiated cells GG-NER via XPC is responsible for repairing a minor fraction of lesions in transcribed genes. UV-C irradiation mainly induces (6-4) photoproducts (6-4PPs) and cyclobutane pyrimidine dimers (CPDs). These lesions are rapidly removed with equal efficiency by TC-NER (43), but CPDs are much less efficiently recognized and slower repaired by GG-NER than by TC-NER (5, 44). The small fraction of XPC-dependent repair therefore likely reflects GG-NER of 6-4PPs rather than CPDs. Strikingly, using a moderate dose of 6 J/m^2^ UV-C irradiation, we observed a complete EGFP-FKBP^F36V^ transcription inhibition in NER deficient cells. This UV dose is estimated to produce on average less than 1 lesion per 10000 bp (43), suggesting that our EGFP-FKBP^F36V^ transgene of 1122 bp is not damaged in each cell. It is therefore not likely that Pol II is physically blocked by DNA damage in the EGFP-FKBP^F36V^ transgene in every cell measured. Indeed, transcription shutdown in response to DNA damage does not only occur *in cis* due to direct physical blockage of elongating Pol II in a particular gene, but also takes place *in trans*, i.e. through genome-wide inhibition of Pol II-mediated transcription initiation and elongation (45–49). Therefore, the lack of EGFP protein synthesis after this moderate UV irradiation of cells with depletion of XPF and CSB most likely reflects the genome-wide shutdown of transcription due to DNA damage induction in a subset of genes. Because of this, it is possible to use the translation of even a relatively short gene like the EGFP-FKBP^F36V^ transgene to monitor DNA damage-induced transcriptional responses and TC-NER activity. In analogy to the RRS assay, we term our assay therefore ‘Recovery of Protein Synthesis’ (RPS).

### Recovery of protein synthesis monitors TC-NER capacity in real-time in living cells

After observing that the RPS assay can be performed by imaging cells that were fixed at defined time points, we tested if RPS can similarly be monitored in real-time in living cells. We therefore imaged siRNA-treated cells, depleted of EGFP-FKBP^F36V^, for 9 h by live cell confocal microscopy after UV irradiation. This clearly showed that in cells treated with control siRNA, the fluorescent EGFP signal recovered already visibly within 1-2 h and continued to increase for 9 h without reaching plateau levels yet. In contrast, in CSB-depleted cells, hardly any fluorescence signal was observed after 1-2 h and only a low recovery of fluorescence signal was visible at later time points (Figure 2A and B; Supplementary movies 1 and 2). These results indicate that measuring RPS in living cells reflects real-time TC-NER activity. In future studies, it might therefore be possible to use the RPS assay to measure and compare TC-NER activity between individual cells, which could for instance be combined with single cell sequencing or proteomics techniques to determine if any observed differences have a molecular (epi)genetic and/or proteomic basis (50).

**Figure 2.**
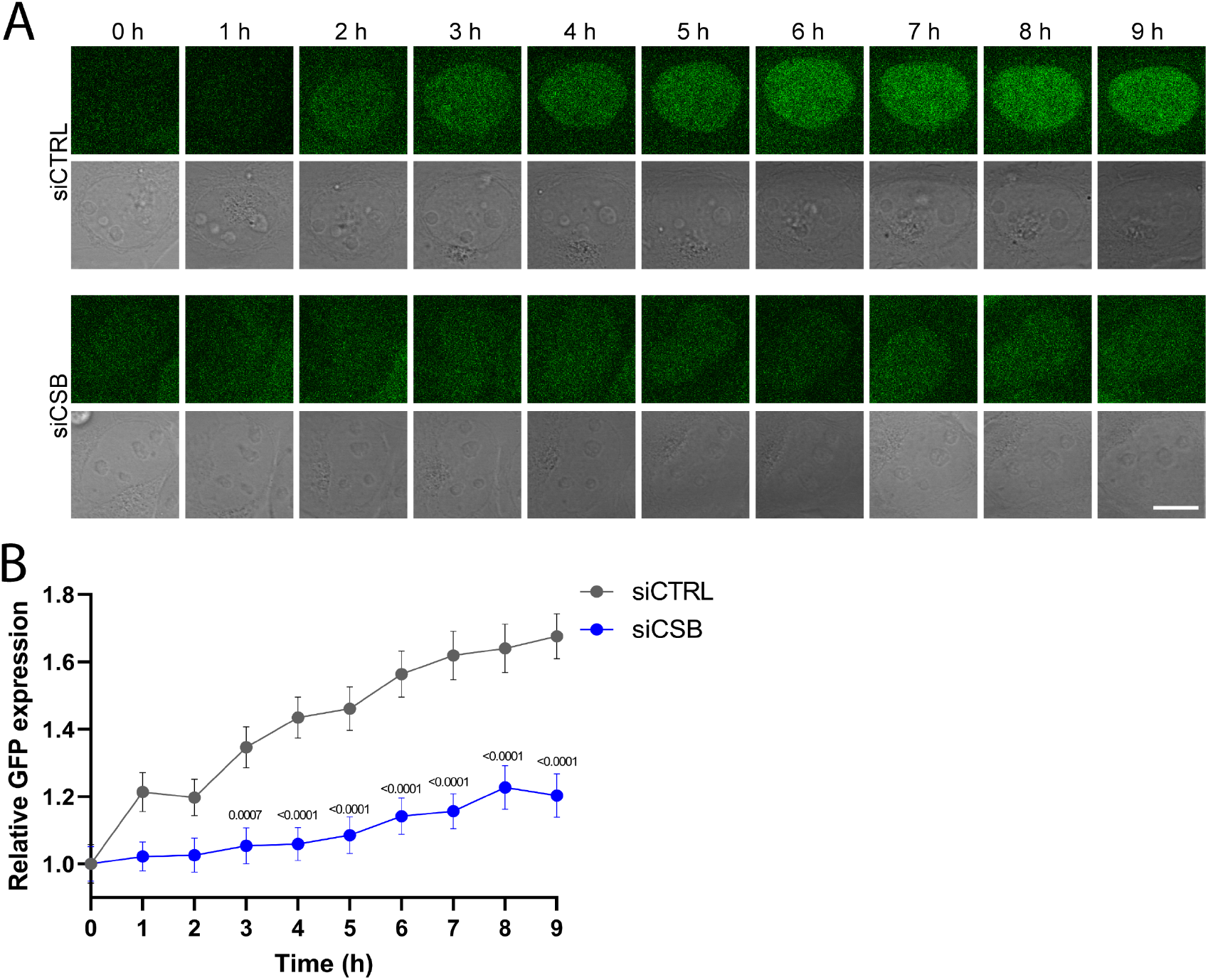
Recovery of protein synthesis monitors TC-NER in living cells. (**A**) Representative live cell imaging pictures of EGFP-FKBP^F36V^-expressing U2OS cells transfected with control (siCtrl) or CSB (siCSB) siRNA. Cells were incubated with dTAG13 for 8 h, irradiated with 6 J/m^2^ UV-C and imaged every hour as indicated. Scale bars, 10 µm (**B**) Quantification of GFP signal of EGFP-FKBP^F36V^-expressing U2OS cells treated and imaged as explained in (**A**). Mean and S.E.M. of at least 89 cells from two independent experiments. Statistical difference determined with unpaired t-test is indicated.

### Recovery of protein synthesis can be performed with chemicals that affect TC-NER

To determine if RPS accurately reflects TC-NER activity in response to DNA damaging agents other than UV irradiation, we exposed siRNA-treated and EGFP-depleted cells to cisplatin for 2 h (Figure 3A). Cisplatin is a commonly used chemotherapeutic drug that mainly creates 1,2-d(GpG) and 1,2-d(ApG) intrastrand crosslinks that effectively inhibit transcription and are repaired via TC-NER (51–54). Similar as after UV irradiation, we observed that the EGFP fluorescence signal in control siRNA-treated cells recovered after cisplatin exposure (Figure 3B and C). In contrast, no fluorescence recovery was observed at all in CSB-depleted cells after cisplatin exposure. These results illustrate the versatility of the RPS assay to measure TC-NER and transcription restart activity in response to different types of genotoxic insults.

**Figure 3.**
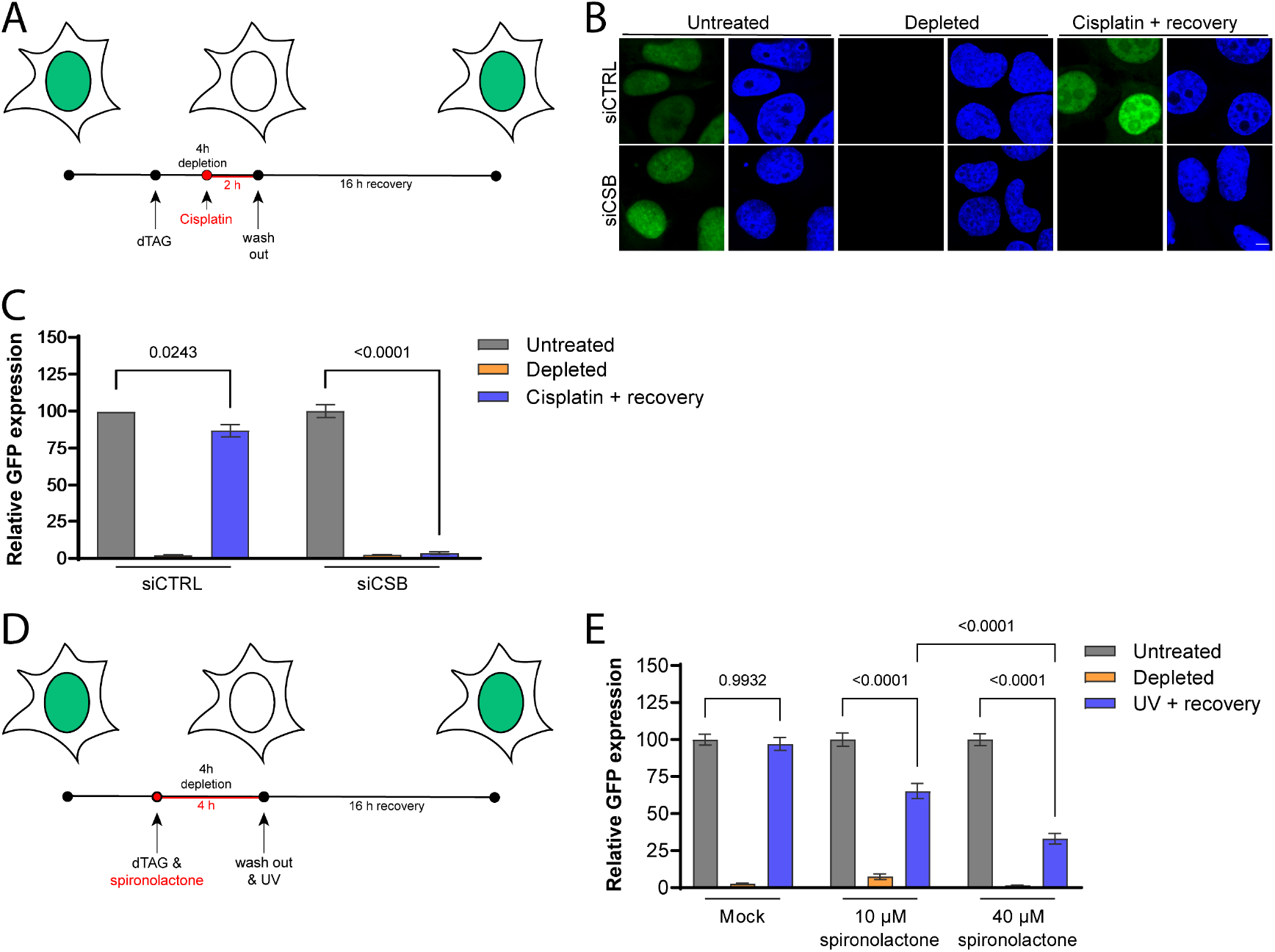
Recovery of protein synthesis after cisplatin or chemical NER inhibition. (**A**) Timing scheme of the Recovery of Protein Synthesis assay using cisplatin. (**B**) Representative images of fixed EGFP-FKBP^F36V^-expressing U2OS cells transfected with control (siCtrl) or CSB (siCSB) siRNA that were either untreated, incubated with dTAG13 for 4 h (‘depleted’) or with dTAG13 for 4 h and 100 µM cisplatin for 2 h and left to recover for 16 h (‘Cisplatin + recovery’). Scale bars, 5 µm (**C**) Quantification of GFP expression levels from siRNA-treated cells imaged and treated as explained in (**B**). Mean and S.E.M. of three independent experiments. (**D**) Timing scheme of the Recovery of Protein Synthesis assay using spironolactone. (**E**) Quantification of GFP expression levels from mock or spironolactone-treated EGFP-FKBP^F36V^-expressing U2OS cells that were either untreated, incubated with dTAG13 for 4 h (‘depleted’) or with dTAG13 for 4 h and then irradiated with 6 J/m^2^ UV-C and left to recover for 16 h (‘UV + recovery’). Mean and S.E.M. of three independent experiments. Statistical difference determined with One way ANOVA is indicated.

We subsequently tested if the RPS assay is compatible with pharmacological inhibition of NER (Figure 3D). We exposed EGFP-depleted cells for 4 h to spironolactone, which induces rapid and reversible degradation of TFIIH subunit XPB (Supplementary Figure 2A) and interferes with NER (55). Indeed, increasing concentrations of spironolactone led to a correspondingly stronger RPS defect in UV irradiated cells (Figure 3E and Supplementary Figure 2B). This shows that the RPS assay can be effectively used to monitor the impact of chemical inhibitors on TC-NER activity. Together, these results indicate that the RPS assay is likely very suited for screening purposes. RPS could be used to screen for genotoxic compounds that block transcription and require TC-NER to overcome the transcription blockage, like cisplatin. Moreover, this type of screening could be combined with high-content screening approaches using genomic siRNA, shRNA or sgRNA libraries to identify the factors involved in repair of these lesions. Additionally, RPS could be used in high-content screening approaches, using fixed or living cells, to screen for compounds that inhibit transcription-coupled DNA repair.

### RPS can be used as diagnostic tool for TC-NER activity with any degradable protein

After establishing the RPS assay by monitoring novel synthesis of dTAG-mediated degraded EGFP in U2OS cells, we tested if instead also other proteins and/or other cell types, including patient-derived fibroblasts, can be used for RPS. We therefore first attempted RPS by monitoring expression of the androgen receptor (AR) after its depletion with Bavdegalutamide (ARV-110) (Figure 4A). ARV-110 is a PROTAC degrader that induces AR ubiquitination and subsequent degradation (56). Imaging of AR protein levels in human hepatoma Hep3B cells stably overexpressing AR fused to EGFP (34, 35) showed that EGFP-AR protein levels were already slightly reduced 16 h after UV irradiation in CSB-depleted cells without the addition of ARV-110 to cells. Even more so, we observed that after ARV-110 addition, the EGFP-AR protein levels only clearly recovered in cells treated with control siRNA, but not in cells treated with siCSB (Figure 4B).

**Figure 4.**
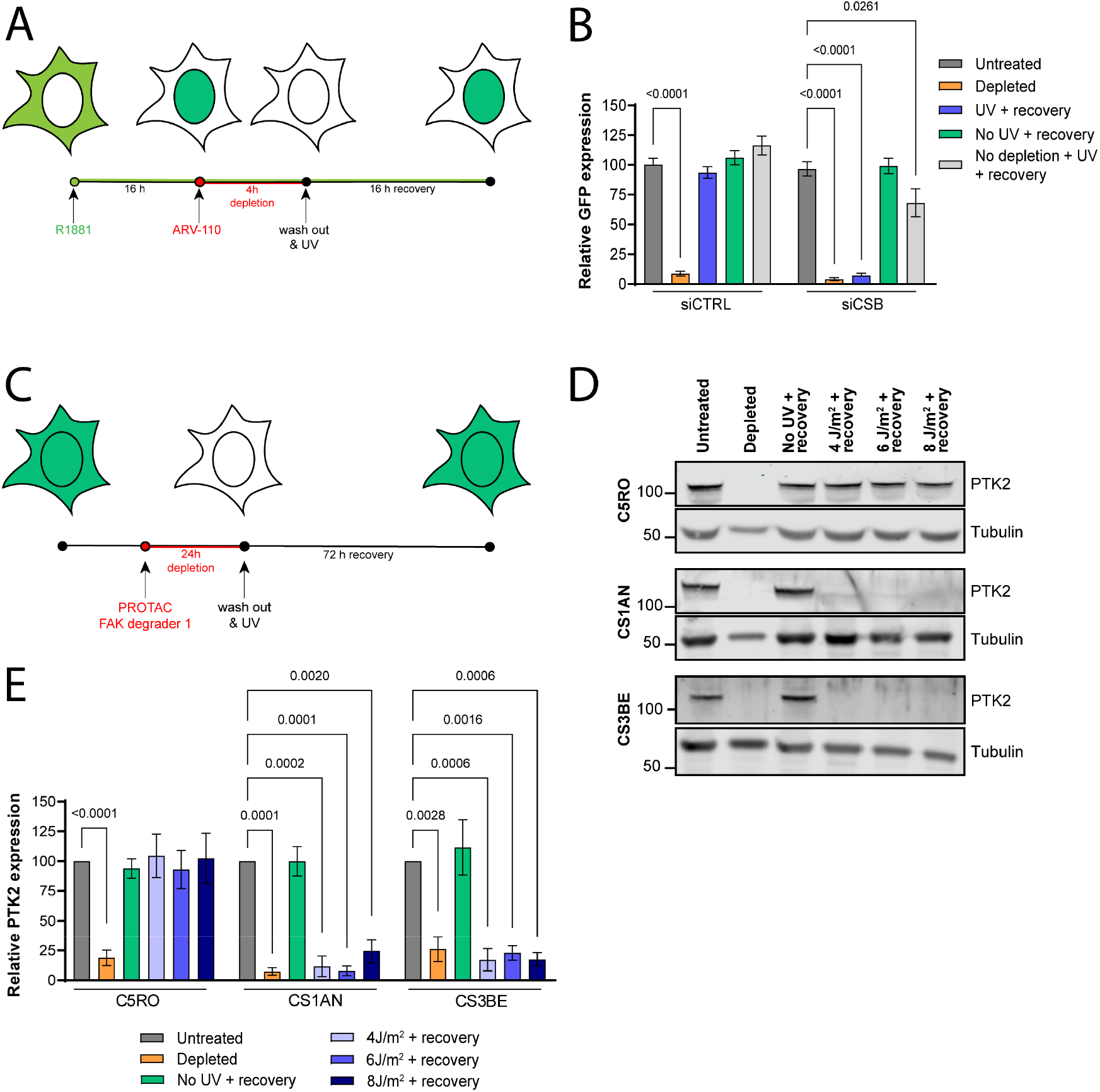
Novel synthesis of PROTAC-degraded proteins to monitor TC-NER. (**A**) Timing scheme of the Recovery of Protein Synthesis assay using ARV-110. (**B**) Quantification of GFP fluorescence levels in EGFP-AR-expressing Hep3B cells that were either untreated or incubated with 100 nM ARV-110 for 4 h (‘depleted’), incubated with 100 nM ARV-110 for 4 h and then irradiated with 6 J/m^2^ UV-C and left to recover for 16 h (‘UV + recovery’), incubated with 100 nM M ARV-110 for 4 h and left to recover for 16 h (‘no UV + recovery’) or irradiated with 6 J/m^2^ UV-C and left to recover for 16 h (‘No depletion + UV + recovery’). (**C**) Timing scheme of the Recovery of Protein Synthesis assay using PROTAC FAK degrader 1. (**D**) Immunoblot analysis of cell lysate from SV40-immortalized C5RO, CS1AN and CS3BE cells that were either untreated or incubated with 250 nM PROTAC FAK degrader 1 for 24 h (‘depleted’), or incubated with 250 nM PROTAC FAK degrader 1 for 24 h and irradiated with the indicated UV dose and left to recover for 72 h (‘indicated UV dose + recovery’). Immunoblots are stained with antibodies against PTK2 and Tubulin (as loading control) (**E**) Quantification of protein levels based on immunoblots as shown in (**D**). Mean and S.E.M. of four (C5RO) or three (CS1AN and CS3BE) independent experiments. Statistical difference determined with One way ANOVA is indicated.

We subsequently tested if we could similarly use this PROTAC-mediated degradation of endogenously-expressed non-tagged proteins to perform RPS in human fibroblasts of CS patients, using immunoblotting instead of fluorescence imaging. As the AR protein is not clearly expressed in fibroblasts (Supplemental Figure 2C), we instead monitored RPS of the PTK2/FAK protein (Figure 4C), which can be efficiently degraded by exposure to the PROTAC FAK degrader 1 (57). We depleted PTK2 in TC-NER proficient control C5RO fibroblasts and in fibroblasts derived from CSB-deficient (CS1AN) and CSA-deficient (CS3BE) CS patients (58). Immunoblotting of cell lysates from these fibroblasts after UV-irradiation showed that PTK2 protein synthesis only clearly recovered in the TC-NER proficient control cells (Figure 4D and E). These results indicate that monitoring RPS of any protein that can be inducibly depleted or degraded, in any cell type of choice, can be used to monitor TC-NER and/or transcription recovery after DNA damage induction. Moreover, the successful application of RPS in patient fibroblasts indicates that this assay could be useful for clinical diagnosis of CS, as versatile alternative to RRS. As we noticed that resynthesis of PTK2 requires substantially longer time than resynthesis of ectopically expressed AR, it will be useful to test and compare additional PROTACs and their protein targets to determine which are most efficient and cost-effective for potential future (clinical) applications of the RPS assay.

### RPS monitors TC-NER activity in young and old tissues of living organisms

To test if the RPS assay can be applied *in vivo*, in differentiated cells of a living organism, we tested if novel protein synthesis in UV irradiated muscle cells of *C. elegans* depends on TC-NER activity (Figure 5A). NER is well-conserved in *C. elegans* and particularly TC-NER is active in differentiated somatic tissues (59, 60). XPC-1 and CSB-1 are *C. elegans* orthologs of XPC and CSB, respectively. We generated transgenic wild type and XPC-1- and CSB-1-deficient animals expressing GFP fused to an auxin-inducible degradation tag (AID::GFP) and *Arabidopsis* TIR1 (fused to mRuby) (39) under control of the *eft-3* promotor driving ubiquitous expression including in muscle cells. TIR1 forms an E3 ubiquitin ligase complex that can be activated by culturing animals on the auxin plant hormone indole-3-acetic acid, which leads to ubiquitylation of the AID tag and subsequent proteasomal degradation of AID::GFP in body wall muscle cells (Figure 5B). 48 h after UV irradiation of *C. elegans* cultured on auxin for 2 h (Figure 5A), we imaged living animals by confocal microscopy and observed that in wild type and XPC-deficient animals the AID::GFP fluorescence had returned to the same level as that of animals not treated with auxin (Figure 5B and 5C). Contrarily, AID::GFP fluorescence levels in UV-irradiated CSB-1-deficient animals remained strongly reduced. Previous survival experiments had suggested that XPC-1 can partially compensate for the lack of repair in active genes in somatic cells of TC-NER-deficient *C. elegans* (40, 61). In line with this, we observed that additional loss of XPC-1 in CSB-1-deficient animals further reduced AID::GFP fluorescence levels after UV. These results confirm that XPC-1 acts partially redundant to CSB-1, as was also observed in human cells (Figure 1E and F).

**Figure 5.**
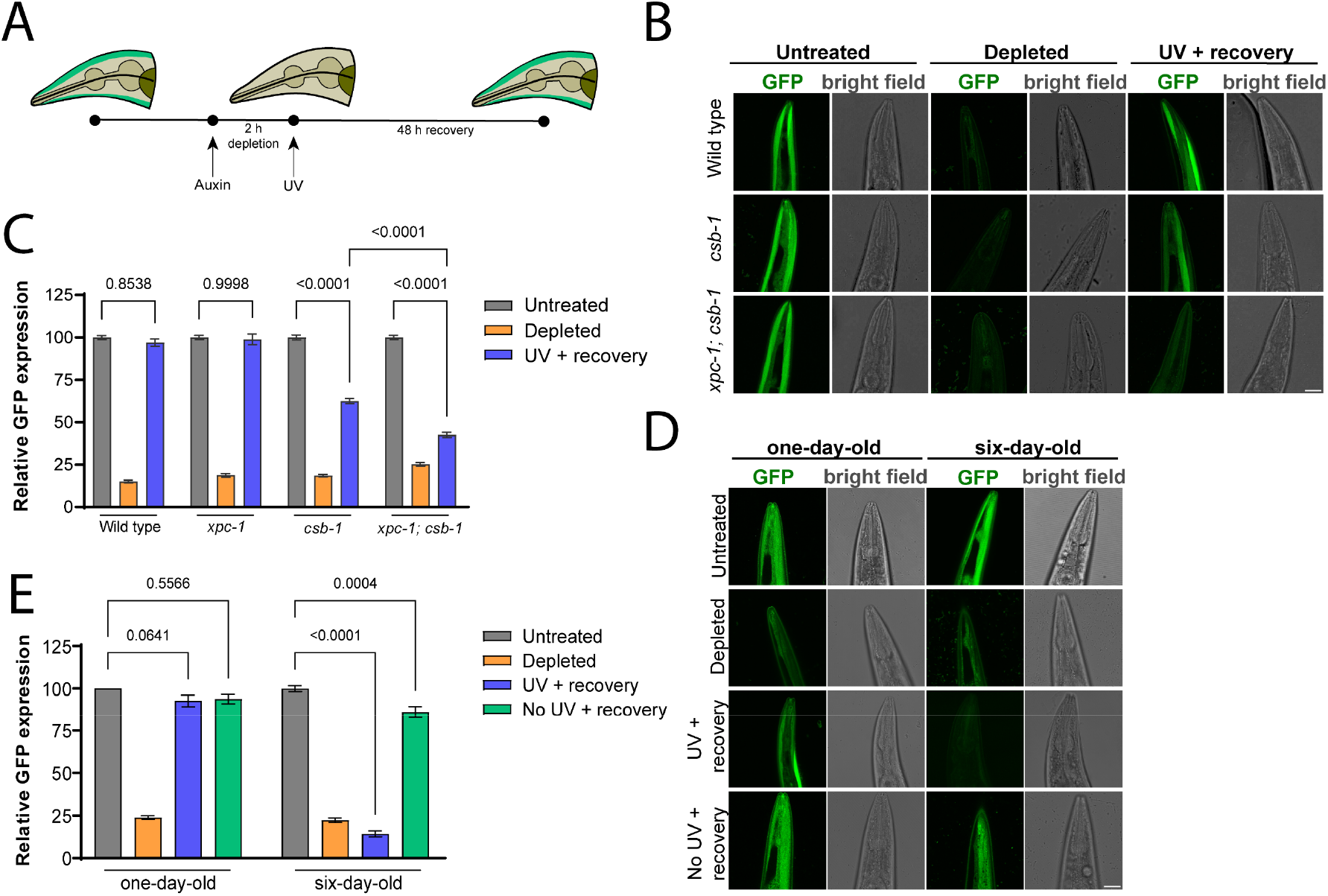
Recovery of protein synthesis assay in muscle cells of *C. elegans*. (**A**) Timing scheme of the Recovery of Protein Synthesis assay using UV-irradiated *C. elegans*. Depicted is a schematic drawing of the head of *C. elegans* with body wall muscle cells expressing green colored GFP. (**B**) Representative confocal images of living wild type, *csb-1* and *xpc-1; csb-1* animals expressing AID::GFP (and TIR1::m-Ruby, not depicted) under control of the *eft-3* promoter in body wall muscles, shown here in the head of *C. elegans*. Animals were either untreated, cultured on 100 mM auxin for 2 h (‘depleted’) or cultured on 100 mM auxin for 2 h and then irradiated with 120 J/m^2^ UV-B and left to recover for 48 h (‘UV + recovery’). Scale bar, 25 mm. (**C**) Quantification of GFP fluorescence levels in muscle cells of wild type, *xpc-1, csb-1* and *xpc-1; csb-1* animals expressing AID::GFP and treated and imaged as explained in (**B**). Mean and S.E.M. of three independent experiments. (**D**) Representative confocal images of living one-day old and six-day old wild type animals expressing AID::GFP in body wall muscles, shown here in the head of *C. elegans*. Animals were either untreated, cultured on 100 mM auxin for 2 h (‘depleted’), cultured on 100 mM auxin for 2 h and then irradiated with 120 J/m^2^ UV-B and left to recover for 48 h (UV + recovery) or cultured on 100 mM auxin for 2 h and left to recover for 48 h (‘no UV + recovery’). Scale bar, 25 mm. (**E**) Quantification of GFP fluorescence levels in muscle cells of one-day-old and six-day-old wild type animals expressing AID::GFP and treated and imaged as explained in (**D**). Mean and S.E.M. of three independent experiments. Statistical difference determined with One way ANOVA is indicated.

As the RPS assay accurately reflects the repair of DNA damage in active genes in muscle cells, we wondered whether it can be used to reveal changes in DNA repair activity when animals grow older, as DNA repair is proposed to decline with age (62). We therefore compared AID::GFP protein levels after UV irradiation in one-day-old wild type adult animals to those in six-day old animals. Whereas UV-irradiated one-day-old animals and unirradiated one-day-old and six-day-old animals all fully recovered GFP fluorescence levels after culturing on auxin, strikingly, UV-irradiated six-day-old animals showed no fluorescence recovery at all (Figure 5D and E). These results suggest that aging *C. elegans* lose their ability to repair DNA damage in actively transcribed genes, at least in muscle cells. These results are in line with previously observed reduced repair in six-day-old animals using gene-specific qPCR assays (63). Together, our results show that the RPS assay can be applied in living, multicellular organisms to compare transcription-coupled DNA repair activity between tissues, different genetic backgrounds and/or between developmental stages or during aging. It will be interesting to investigate if a similar reduction of TC-NER capacity can be measured in aging tissues of e.g. mouse models of aging, as accumulating DNA damage leading to transcription stress is considered one of the driving forces of the aging process (22, 64).

### Prospects and limitations of the RPS assay

Several proteins have been implicated in transcription restart following DNA damage induction and DNA repair, rather than DNA repair itself (65–68). It is important to note that the RPS assay, although versatile in its applicability with different cell types both *in vitro* and *in vivo*, cannot be used to distinguish between TC-NER and restart defects. To make such a distinction, results should be compared to those obtained in assays directly measuring TC-NER such the amplified UDS (30) or strand-specific ChIP-Seq (31) assays. Also, it is unknown if with the DNA damage doses that we used translation itself is affected. Previous reports have shown inhibition of protein synthesis after DNA damage induction, but this occurred only with relatively high UV doses (69–71). Possibly, our assay could be useful to identify factors involved in regulating the translational response to DNA damage as well. As the RPS assay is easily adjustable for use with different cell types, degradable target proteins and/or experimental readouts, it is expected that the assay will be very useful for future screening purposes utilizing living cells or even tissue models such as organoids or various model organisms.

## DATA AVAILABILITY

Plasmids, cell lines and *C. elegans* strains generated in this study are available upon reasonable request.

## SUPPLEMENTARY DATA

Supplementary Data are available at NAR online.

## ACKNOWLEDGEMENT

We thank Gert van Cappellen of the Erasmus MC Optical Imaging Center for microscope support. Some *C. elegans* strains were provided by the Caenorhabditis Genetics Center (funded by NIH Office of Research Infrastructure Programs P40 OD010440) and the National Bioresource Project for the nematode.

