## supplemental figures for "Recovery of protein synthesis to measure transcription-coupled DNA repair in living cells and tissues"

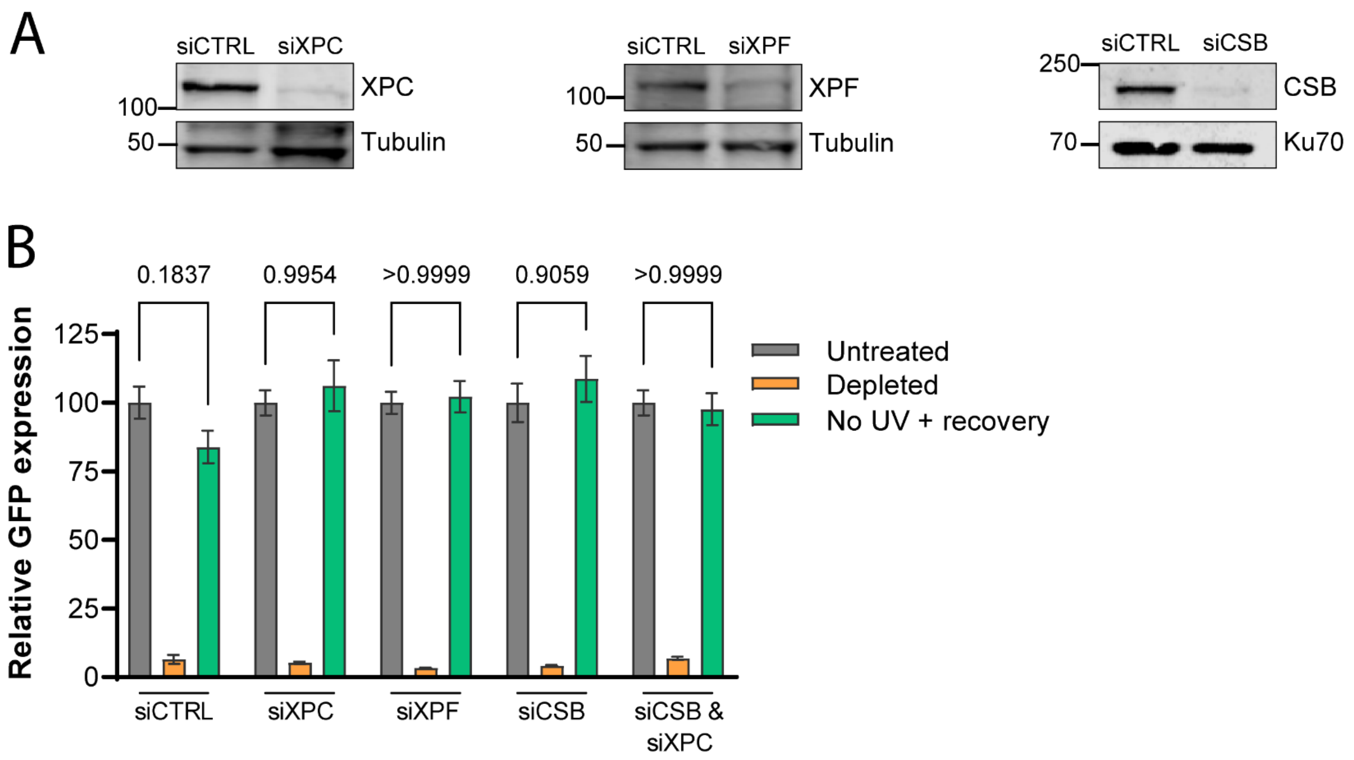


**Supplementary Figure 1.** Recovery of protein synthesis without UV irradiation. (**A**) Immunoblots of total U2OS cell lysates demonstrating the efficiency of siRNA-mediated protein depletion with siRNAs against XPC, XPF and CSB, probed with antibodies against the respective proteins and against tubulin or Ku70 as loading control. (**B**) Quantification of GFP expression levels in fixed EGFP-FKBP$^{F36V}$-expressing U2OS cells transfected with control, XPC, XPF, CSB or CSB and XPC siRNA that were either untreated, incubated with dTAG13 for 4 h ('depleted') or incubated with dTAG13 and left to recover for 16 h ('No UV + recovery'). Mean and S.E.M. of two independent experiments. Statistical difference determined with One way ANOVA is indicated.

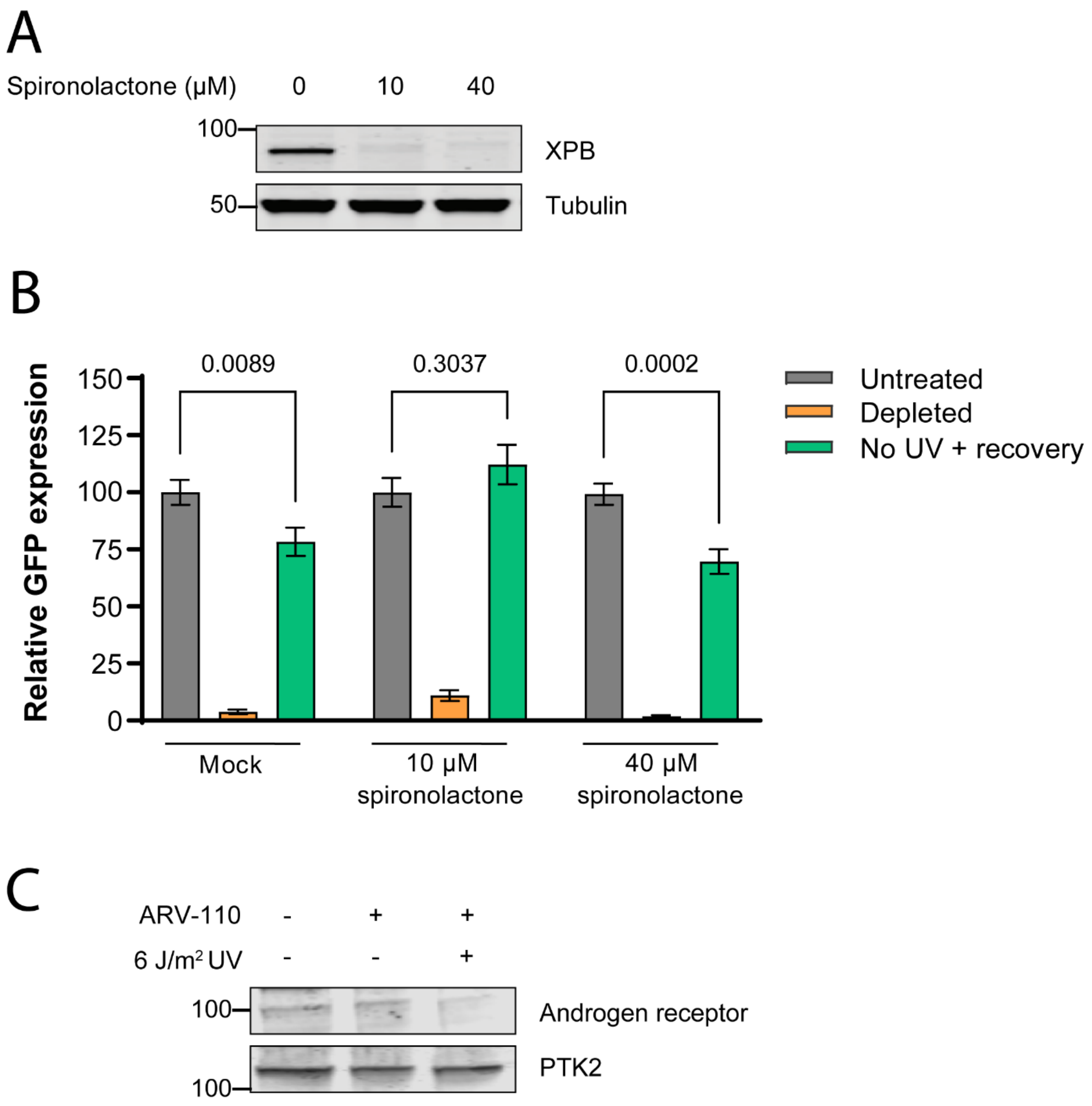


**Supplementary Figure 2.** Spironolactone inhibition of XPB and PTK2 expression in human fibroblasts. (**A**) Immunoblot analysis showing XPB protein levels in EGFP-$\text{FKBP}^{F36V}$-expressing U2OS cells treated with the indicated spironolactone concentrations for 4 h. Immunoblot is stained with antibodies against XPB and tubulin as loading control. (**B**) Quantification of GFP expression levels from mock or spironolactone-treated EGFP-$\text{FKBP}^{F36V}$-expressing U2OS cells that were either untreated, incubated with dTAG13 for 4 h ('depleted') or with dTAG13 for 4 h and then irradiated with 6 $J/m^2$ UV-C and left to recover for 16 h ('UV + recovery') or with dTAG13 for 4 h and left to recover for 16 h ('no UV + recovery'). Mean and S.E.M. of two independent experiments. Statistical difference determined with One way ANOVA is indicated. (**C**) Immunoblot analysis of total cell lysate of C5RO cells either mock treated, or treated with 100 nM ARV-110 for 4 h and 6 $J/m^2$ UV irradiation, as indicated, using antibodies against the Androgen receptor and PTK2.

**Supplementary movie S1**

Movie showing recovery of GFP protein synthesis, reflecting TC-NER activity, in living U2OS cells expressing EGFP-FKBP$^{F36V}$ and transfected with control siRNA. Cells were incubated with dTAG13 for 8 h, irradiated with 6 J/m$^2$ UV-C and immediately imaged every 12 min. Upper panel shows GFP fluorescence and lower panel shows bright field image. Related to Figure 2.

**Supplementary movie S2**

Movie showing recovery of GFP protein synthesis, reflecting TC-NER activity, in living U2OS cells expressing EGFP-FKBP$^{F36V}$ and transfected with CSB siRNA. Cells were incubated with dTAG13 for 8 h, irradiated with 6 J/m$^2$ UV-C and immediately imaged every 12 min. Upper panel shows GFP fluorescence and lower panel shows bright field image. Related to Figure 2.
